# Exosomes purified from a single cell type have diverse morphology and composition

**DOI:** 10.1101/094045

**Authors:** Davide Zabeo, Aleksander Cvjetkovic, Cecilia Lässer, Martin Schorb, Jan Lötvall, Johanna Höög

## Abstract

Extracellular vesicles (EVs) are produced by all known organisms and have important roles in cell communication and physiology. Exosomes are known in the literature to be small round EVs (40 to 100 nm in diameter) and are commonly purified with a serial ultracentrifugation protocol followed by density gradient floatation. Great morphological diversity has been described before regarding EVs found in body fluids such as blood plasma, breast milk and ejaculate. However, a detailed morphological analysis has never been performed on exosomes purified from a single cell type.

Therefore, the aim of this study was to analyze and quantify via multiple electron microscopy techniques the morphology of exosomes purified from the human mast cell line HMC-1. The results revealed a novel spectrum of diversity in exosomes, which suggests that subpopulations of exosomes with different and specific functions might also exist. Our findings therefore argue that a new and more efficient way of defining exosome subpopulations is necessary. A system was proposed where exosomes were classified into nine different categories according to their size and shape. Three additional morphological features could also be found in exosomes regardless of their classification.

These findings show that morphological diversity is found among exosomes purified from a single cell line, similarly to what was previously observed for EVs in body fluids. This knowledge can help improving the interpretation of experimental results and widening our general understanding of the biological functions of exosomes.

## Introduction

Communication is critical for survival in the natural environment. On a cellular level, cell communication can be mediated by extracellular vesicles (EVs). All studied organisms from bacteria (1, 2) to animals (3) and plants (4, 5) are known to produce EVs. EVs carry proteins and nucleic acids and deliver them to target cells which may undergo substantial physiological changes as a consequence (6-10). EVs have been identified in several human body fluids, such as blood (11, 12), breast milk (13, 14) and ejaculate (15–17), and also in human tissues, including tumors (18, 19). Thus, understanding EV-mediated communication is important for basic science and medicine alike. Depending on their biogenesis EVs are classified as: apoptotic bodies, which are released by cells undergoing apoptosis, microvesicles, which are shedded directly from the plasma membrane, or exosomes, which are produced in organelles called multivesicular bodies (MVBs)(3, 20, 21). MVBs mature from endosomes as their membrane buds inwards in their lumen to produce vesicles (22), which are then called exosomes if released into the extracellular environment (23, 24). However, no purification methods available can separate EVs based on their biogenesis. Instead, vesicles isolated using ultracentrifugation, the most common exosome purification technique (25, 26), followed by floatation in a density gradient, are commonly called exosomes (18, 27). We will also use this definition of exosome in this paper.

Once purified, exosomes are often characterized using negative stain electron microscopy, a method where the surface structure is revealed by coating the vesicles with heavy metal salts (28). Exosomes are commonly considered to be a homogeneously shaped group of vesicles, sized 40-100 nm in diameter (20). On the other hand, an abundance of morphological diversity has emerged when extracellular vesicles from bodily fluids (11–13,15, 16) and cell culture supernatant (29–31) were studied using cryo-electron microscopy (cryo-EM), a method which allows visualization of membrane bilayers and internal features (11–13, 31). However, thorough morphological studies using cryo-EM on exosomes purified from a single cell type are lacking to date. Therefore the purpose of this work was to characterize, in detail, the morphological diversity of exosomes purified from a single cell line (the human mast cell line HMC-1), with the aim of determining if a single cell type can produce multiple types of exosomes.

## Materials and Methods

### Cell culture

The human mast cell line (HMC-1) was cultured in Iscove’s modified Dulbecco medium (IMDM) supplemented with 10% exosome depleted fetal bovine serum, 100 units/ml streptomycin, 100 units/ml penicillin, 2mM L-glutamine and 1.2 U/ml alpha-thioglycerol in incubators kept at 37°C and 5% CO_2_ (all supplements from Sigma, St Louis, MO, USA). FBS was depleted of vesicles by an 18-hours-long centrifugation at 118 000 × g_avg_ (Beckman Coulter Type 45 Ti rotor, k-factor 178.6) at 4°C followed by filtration through a 0.22 μm filter (32).

### Exosome isolation

Exosomes were harvested from HMC-1 cells cultured for three days and a cell viability of at least 98% was ensured by trypan blue staining. EVs were collected by differential centrifugation followed by floatation on a density gradient. Media was collected from cell cultures and spun at 300 × g for 10 minutes to deplete it of cells. The supernatant was carried over to ultracentrifuge tubes (100 ml polypropylene quick-seal tubes, Beckman Coulter) and centrifuged at 16 500 × g_avg_ for 20 minutes at 4°C in order to pellet larger vesicles such as apoptotic bodies and microvesicles (Type 45 Ti, k-factor 1279). The supernatant was again carried over to new ultracentrifuge tubes and centrifuged at 118 000 × g_avg_ for 3.5 hours at 4°C to pellet the exosomes (Type 45 Ti) (33). The supernatant was discarded and the pellets were resuspended in PBS. An isopycnic density gradient centrifugation was performed on the pellets. 1 ml of the resuspended exosome pellet was bottom loaded together with 3 ml of 60% optiprep by mixing these two together in open top ultracentrifuge tubes (13.2 ml Ultra Clear tubes, Beckman Coulter). A step gradient of 1 ml each with the concentrations 35%, 30%, 28%, 26%, 24%, 22% and 20% (except for the 22% of which 2 ml were added) of optiprep containing 0.25 M sucrose, 10mM Tris and 1mM EDTA was overlaid in that order. The gradient was topped off with approximately 200 μ PBS to completely fill the tube. The gradient was then centrifuged at 178 000 × g_avg_ for 16 hours at 4°C (SW 41 Ti, k-factor 144). A clearly visible band at the intersection where the 20% – 22% were loaded, with an approximate density of 1.12 g/ml, was collected. The floated EVs were then washed by diluting them in approximately 100 ml of PBS and centrifuged at 118 000 × g_avg_ for 3.5 hours at 4°C (Type 45 Ti). The pellet was then finally resuspended in PBS to be used for downstream experiments.

### High pressure freezing of HMC-1 cells for electron microscopy

High-pressure freezing and freeze substitution reduce the artifacts created with chemical fixation at room temperature by carrying out the sample preparation procedure at low temperatures (34-37). HMC-1 cells were centrifuged at 600 × g for 4 min to form a pellet. The sample for high pressure freezing was pipetted up from inside this pellet to maximize the cell concentration. Cells were loaded into membrane carriers (150 mm; Leica Microsystems) and high pressure frozen in a Leica EM Pact1 (Leica Microsystems). A short freeze substitution protocol was applied, using 2% uranyl acetate (UA) for one hour at −90°C (36, 38-40). Samples were embedded in K4M resin and sectioned in 70 nm thin sections placed on copper slot grids. Sections were stained using UA and lead citrate.

### Negative stain of isolated exosomes

Negative stain is the fastest way to prepare a sample for electron microscopy. The treatment covers the sample with a layer of heavy metal salts which creates contrast and allows visualization of the sample surface (28). 5 μl of sample were applied to glow-discharged carbon coated formvar 200 mesh grids and incubated for 10 minutes. Washed two times with PBS and then fixed using 2.5% glutaraldehyde for 5 min. Grids were then washed with filtered distilled water and stained using 2% UA in water for 1 min.

### Cryo-electron microscopy

Cryo-EM allows imaging of samples without the addition of any heavy metals or fixatives, which might cause artifacts, with the drawback of yielding a lower contrast. The sample is so rapidly frozen that the water vitrifies forming no ordered crystals and the native structure of the sample is preserved (41, 42). Freshly prepared vesicles were plunge frozen in liquid propane using a Vitrobot (FEI; Eindhoven, The Netherlands). In some instances, 10 nm protein A gold particles were added as fiducial markers (Cell Microscopy Core; Utrecht, The Netherlands).

### Electron microscopy

Thin sections and negative stain samples were imaged on a Leo 912AB Omega TEM operated at 120 kV. Cryo-electron microscopy was performed on either a CM200 (Karolinska Institute, Stockholm) or a Tecnai F30 (EMBL, Heidelberg; (FEI; Eindhoven, The Netherlands)).

### Exosome analysis and classification

Electron micrographs of exosomes were analyzed with the IMOD software (43). Vesicles were included in the analysis only if they presented a visible membrane bilayer. Vesicles were counted and their size was measured (diameter if round, length if elongated).

## Results

### Morphological variability of vesicles inside MVBs in cells

In order to investigate the morphological nature of exosomes produced by the human mast cell line (HMC-1), multiple EM techniques were employed. HMC-1 cells were imaged after high-pressure freezing, freeze substitution and sectioning as reported in Materials and Methods.

It was possible to distinguish several MVBs inside of the cells (Figure 1A; white boxes). The same MVBs are also shown with higher magnification (Figure 1B-D), together with examples of MVBs that were found inside other cells (Figure 1E-H). MVBs contained structurally diverse vesicles (white arrows) as well as normal single vesicles. The same MVB contained vesicles of variable sizes or filled with more electron dense material (Figure 1C-D). Smaller vesicles were seen inside larger vesicles (Figure 1C-F) and an entire MVB was found inside of another, larger MVB together with a number of vesicles (Figure 1E). Many MVBs were surrounded by vesicles that were located in their proximity or in some cases docked onto the outer side of their membrane (Figure 1B, G).

**Figure 1:**
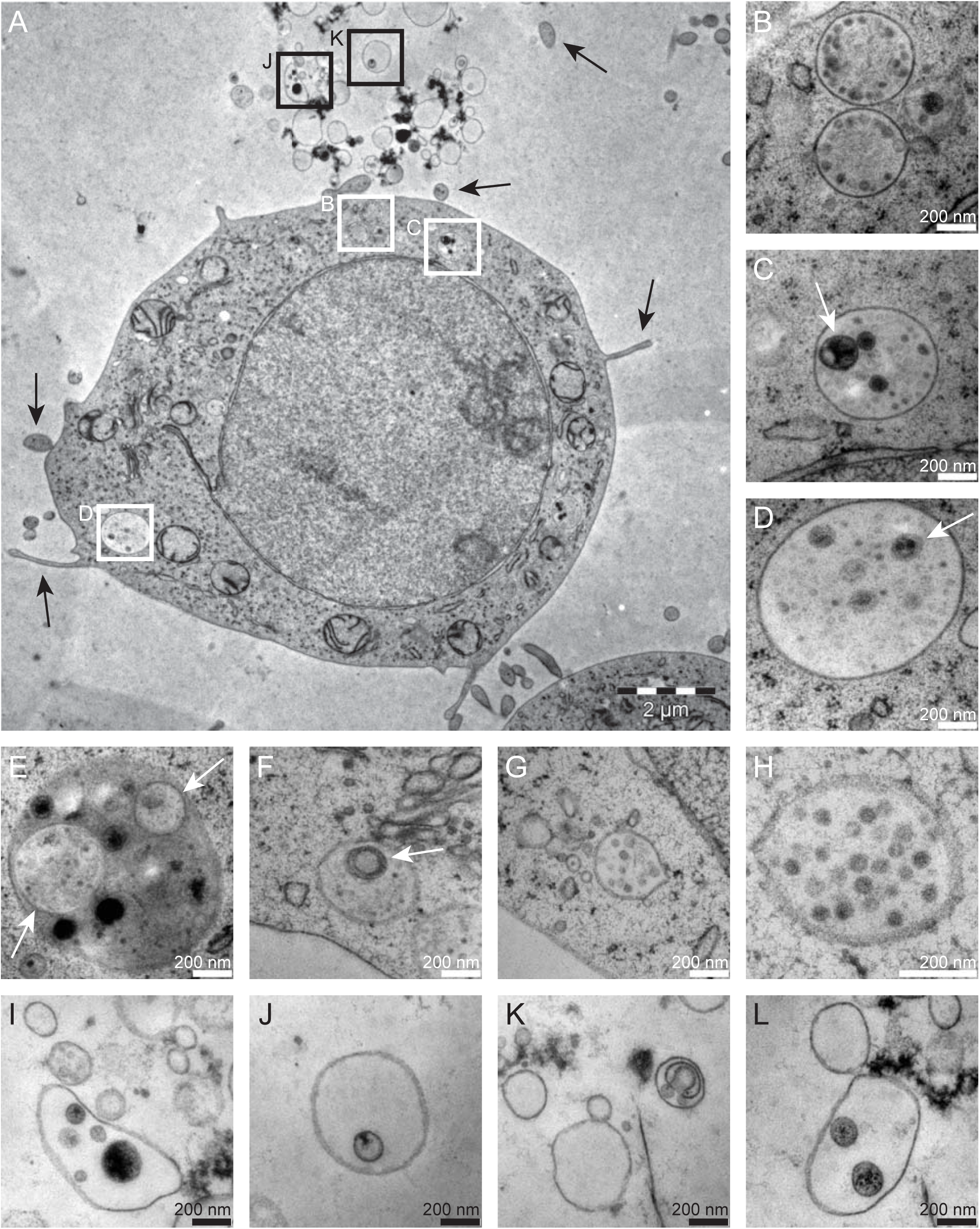
Thin sections show MVBs contained in cultured HMC-1 cells and extracellular vesicles. A) HMC-1 cell containing MVBs (white boxes) and some EVs in proximity of it (black boxes). The black arrows point at cell protrusions. The letters next to the boxes refer to the adjacent higher magnification panels in the figure. B-H) MVBs in higher magnification. White arrows point at vesicles containing one or more smaller vesicles. J-L) EVs in higher magnification.

Figure 1 also highlights some extracellular vesicles present in the cellular milieu (Figure 1A, I-L; black boxes). EVs could be recognized by their transparent and lightly-textured lumen and should not be confused with cross sections of cell protrusions (Figure 1A; black arrows), which had a darker ribosome-filled texture which resembled that of the cytoplasm. Morphological differences were also present in these EVs, regarding shape (Figure 1I-J), electron density and number of smaller vesicles that were contained inside of them, which varied from none to one or two or even more (Figure 1K, J, L and I respectively). A complicated structure was seen in one EV (Figure 1K), showing that non-spherical membranous compartments can be found in the extracellular environment. Although it’s unknown whether the biogenesis of these EVs was MVB-associated and if their variable morphology was linked to the structural properties of exosomes, their existence still implies a level of complexity in EVs moieties that might have been overlooked.

These findings show a morphological variability of extracellular vesicles in cell culture medium and in vesicles inside MVBs, even before their release into the environment. Therefore, their morphological diversity after a classical exosomes isolation procedure was examined.

### Exosomes purified from HMC-1 can be classified into nine categories

Exosomes were isolated using differential ultracentrifugation followed by a density gradient flotation purification. Exosomes were imaged using two different techniques: negative staining and cryo-EM (Figure 2A-B respectively). With negative staining, differences in both shape and size were observed, with a long tubule-like vesicle (red arrow) surrounded by many other round vesicles with variable diameters (green arrows).

**Figure 2:**
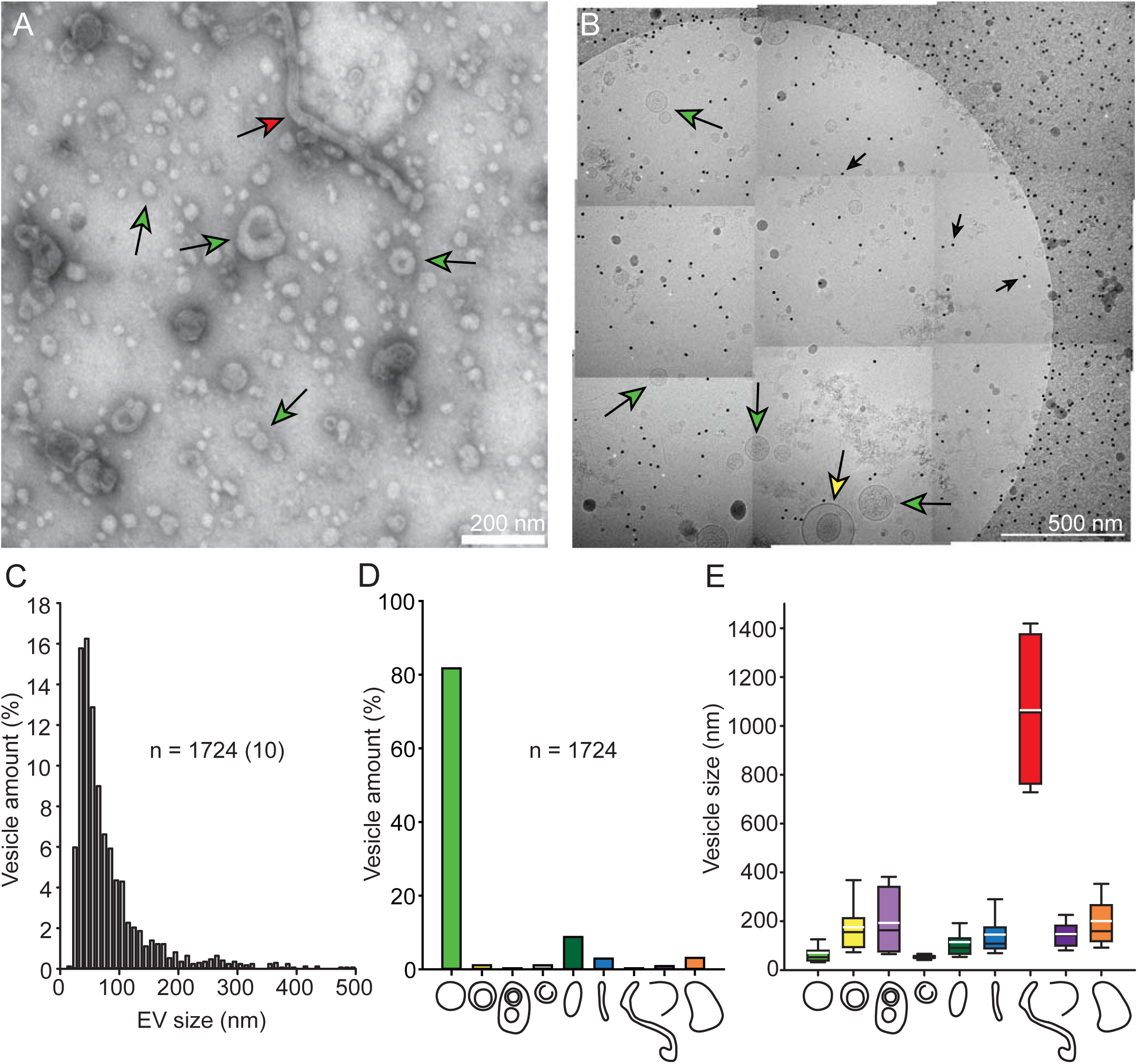
Morphologically diverse exosomes purified from HMC-1. A) Exosomes visualized after negative staining. At least two different categories of vesicles can be identified in this picture: single vesicles and a long tubule (green arrows and red arrow respectively). B) Montage of 9 cryo-electron micrographs. Green arrows point at single vesicles, yellow arrow points at a double vesicle, small black arrows point at fiducial gold markers. The circular shape is the carbon edge of the holey carbon grid. C) Size distribution of all vesicles included in the analysis (n=1724). In brackets is indicated how many vesicles were larger than 500 nm. D) Percentage of total vesicles that belonged to each morphological category. E) Size distribution for each vesicle category. The top and bottom boundaries of the boxes represent the 75th and 25th percentiles. The top and bottom whiskers represent the 90th and 10th percentiles. The black line in the box represents the median while the white line represents the mean.

Cryo-EM yielded additional information on the vesicles regarding their structure, membrane and lumen, since lipid bilayers and vesicle internal structures could be visualized. Figure 2B shows an example of montage of 9 electron micrographs covering the entire hole of a carbon grid. Exosomes were counted in a total of 410 micrographs, their size was measured and they were classified in 9 different categories depending on their morphology, as listed below. In total 1724 exosomes were counted and 75% of them measured between 30 and 100 nm in size, with the rest being mostly larger (Figure 2C). The percentage of vesicles that belonged to each category and their size distribution within each category are shown in Figure 2D-E respectively. Examples of vesicles for every category are found in Figures 3 and 4.

**Figure 3:**
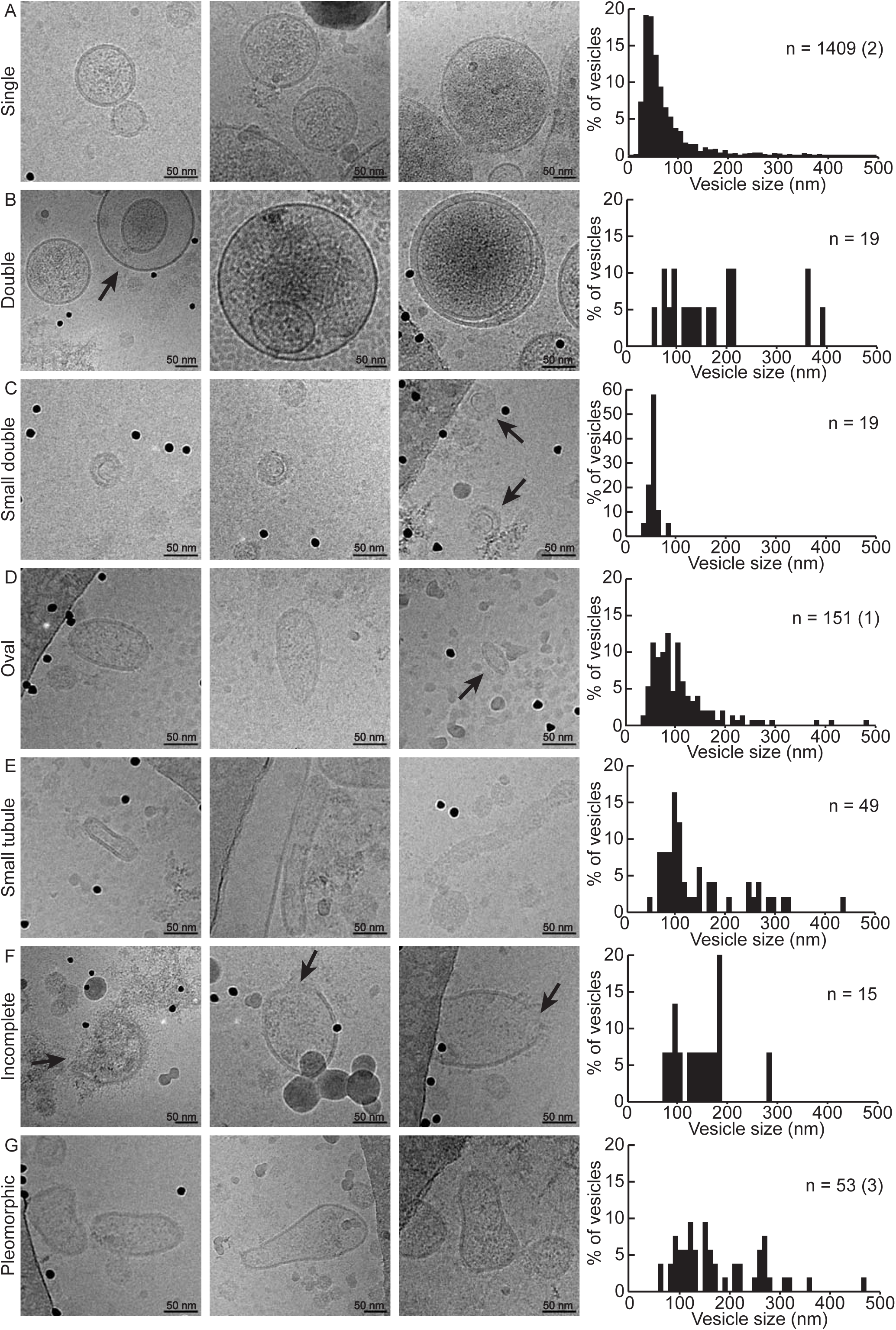
Gallery of exosome categories. A-G) Three example vesicles for each category are shown, followed by the size distribution of the category. Y axis shows the percentage of vesicles within each particular category. Sample size (n) is indicated and in brackets is how many vesicles in that category exceeded 500 nm in size.

**Figure 4:**
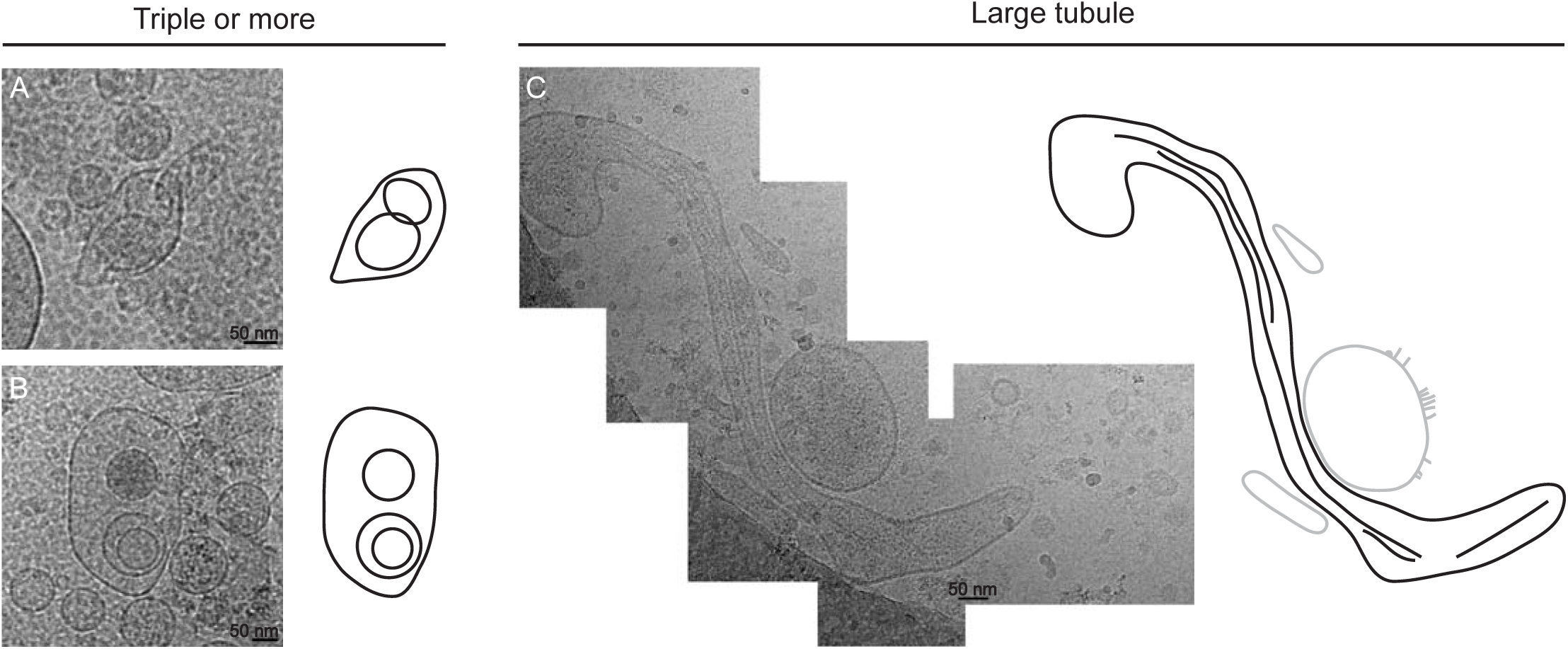
Gallery of exosome categories. A-B) Two examples for the category of triple vesicles or more with drawings to highlight their structure. C) One example of a filamentous large tubule with drawing (large tubule in black, nearby vesicles in gray).

#### Single vesicle (n = 1409)

Single vesicles represented the majority of all counted exosomes (81.7%; Figure 2D). They were delimited by a membrane bilayer and had a round shape. Their size range was 71.22 ± 58.42 nm (mean ± stdev; Figures 2E and 3A).

#### Double vesicle (n *=* 19)

Double vesicles contained a smaller vesicle enclosed into a larger one and their shape was either round or slightly elongated. They were larger in size than the average single vesicle (174.26 ± 100.80 nm; t-test, P < 0.001; Figure 3B).

#### Triple vesicle or more (n *=* 4)

Vesicles fell into this category when two or more vesicles were contained inside a larger one. They were found in different combinations, having three vesicles inside one another, or having two smaller vesicles both inside the same large one (Figure 4A). Only four vesicles were classified to be part of this category, the smallest being 65 nm long and the largest 380 nm (Figure 2E).

#### Small double vesicle (n *=* 19)

These vesicles differ from regular double vesicles since the inner structure was thinner than a bilayer and discontinuous, meaning it did not form a full circle in its shape (Figure 3C).

They were also significantly smaller than double vesicles, having a size range of 54.10 ± 10.46 (*t*-test, P < 0.001; Figure 2E).

#### Oval vesicle (n *=* 151)

Oval vesicles had a single bilayer similarly to single vesicles, but they were elongated in shape, having only two perpendicular symmetry axes (Figure 3D). Their average size was 114.26 ± 85.89 nm (Figure 2E).

#### Small tubule (n *=* 50)

These vesicles were also characterized by a single bilayer and an elongated shape, but they differed from oval vesicles in the fact that, along their elongated dimension, the membranes on opposite sides of their symmetry axis were parallel to each other, causing these vesicles to have a more tubular shape (Figure 3E). Their size averaged at 144.76 ± 86.95 nm (Figure 2E).

#### Large tubule (n *=* 4)

Large tubules were distinct from small tubules only because of their size. Only 4 large tubules were identified and all of them measured longer than 700 nm, up to 1400 nm (Figures 2E, 4B).

#### Incomplete vesicle (n *=* 15)

These vesicles showed an interrupted membrane and were often surrounded by electron dense material (Figure 3F). It is possible that the breakage occurred during sample preparation.

#### Pleomorphic vesicle (n *=* 53)

All vesicles whose shape could not be classified in any of the previous categories were called pleomorphic. Their size was larger than the average single vesicle (200.58 ± 86.95 nm; *t*-test, P < 0.001; Figure 2E). Examples are shown in Figure 3G.

In addition to all these categories, a complicated structure consisting of a broken membrane located adjacently to several single, double and triple round vesicles was observed (Figure S1). It resembled the category of vesicle sacs previously described for ejaculate vesicles (15).

### Three additional features could be found in vesicles from different categories

In addition to their morphological traits, some vesicles possessed up to three features independently of the category they belonged to. Examples for each feature are shown in Figure 5A-I. The number of vesicles that presented features in each category is summarized in Table S1.

**Figure 5:**
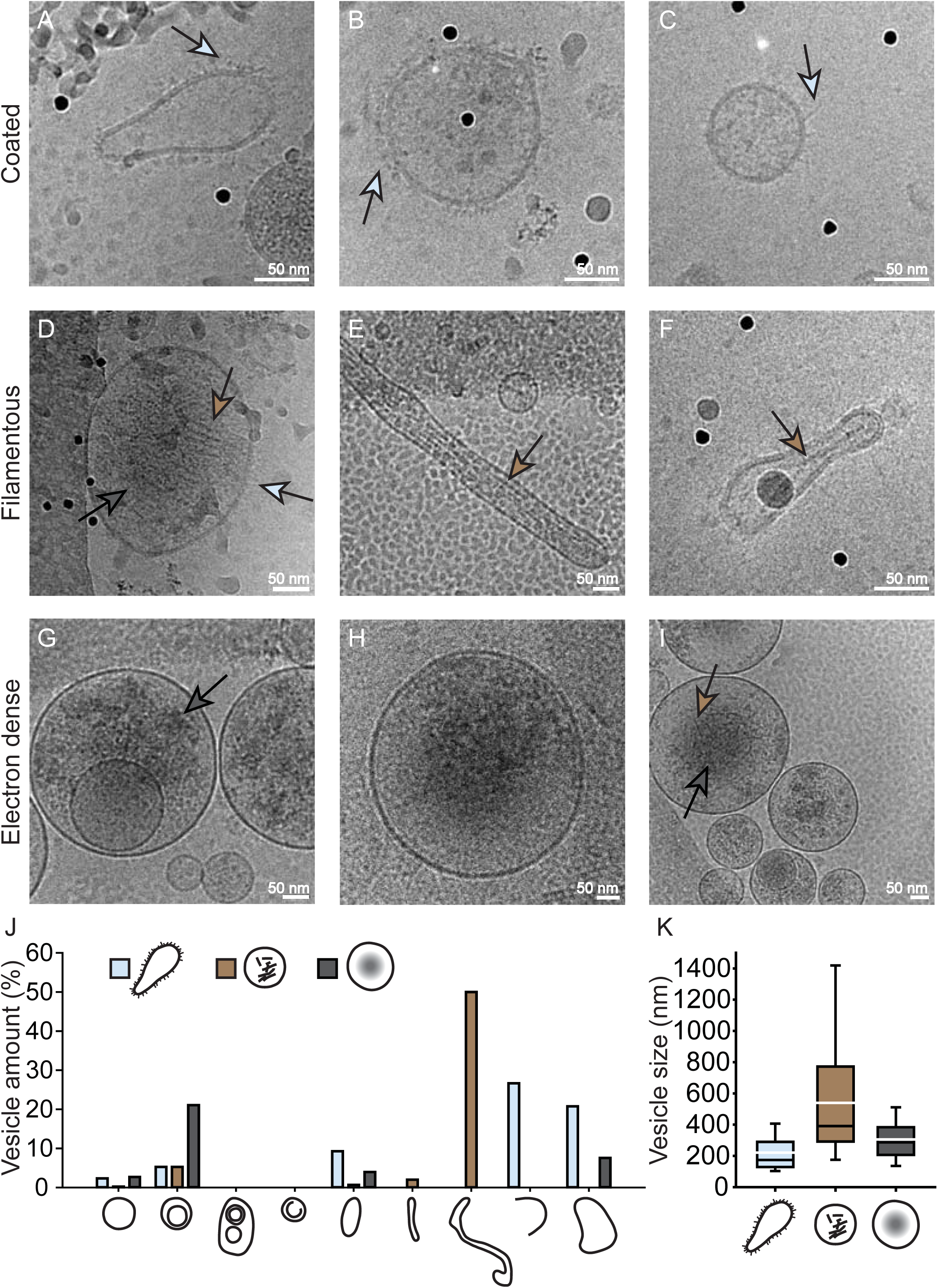
Gallery of exosome features. A-I) Examples of coated (A-D), filamentous (D-F, I) and electron dense vesicles (D, G-I). Light blue arrows point at coating, brown arrows point at filaments and grey arrows point at electron density within vesicles. J) Percentage of each vesicle category that presented each of the features. K) Size distribution of vesicles with features.

#### Coated vesicles

Some vesicles had surface protrusions, electron dense spikes protruding from the membrane, or part of the membrane (Figure 5A-C). Around 3.7% of all vesicles showed this feature (63 out of 1724) and their average size was 220.53 ± 132.96 nm (Figure 5K).

#### Filamentous vesicles

0. 5% of counted vesicles contained filaments in their lumen (Figure 5D-F, I). In vesicles with a tubular shape the filaments were aligned in parallel with the length of the tubule (Figure 5E-F) but in round vesicles filaments could be arranged parallel with each other (Figure 5D) or more randomly distributed (Figure 5I). Their size was very variable due to the fact that filaments were found in single vesicles as well as in small and large tubules (Figure 5K). In fact, 2 out of 4 large tubules were recognized as filamentous (Figure 5J).

#### Electron dense vesicles

These vesicles appeared to be filled with electron dense material, which gave them a darker texture in their lumen compared to nearby vesicles of the same size (Figure 5G-I). Their mean size (305.52 ± 140.79 nm) was above the average size for the total vesicle count (*t*-test, P < 0.001; Figure 5K).

Features were not mutually exclusive, as in some cases a vesicle could present more than a single feature. Figure 5D for example shows a vesicle in which all three features can be identified.

## Discussion

EVs of many different morphologies have previously been reported when studying bodily fluids using cryo-electron microscopy (11–13, 15, 16) and some morphological variability has been mentioned in exosomes from single cell types (29–31). However, thorough quantitative morphological studies on exosomes isolated from a single cell type could not be found to date. Exosomes are commonly known in the literature as small, round EVs of the size of 40 to 100 nm in diameter (20), but our paper revealed with multiple EM techniques that exosomes produced by HMC-1 cells included morphologically diverse subpopulations. It should be noted that exosomes were purified via density gradient floatation and that all analyzed vesicles were collected from a single band of approximately 1.12 g/ml. The morphological variability was therefore observed in a population of exosomes which were produced by a single cell type and which had equal densities.

The morphological variability of the exosomes suggests the existence of different subpopulations of them which possess different functions and biochemistry. For instance, coated vesicles exhibited protrusions on their surface which other vesicles lacked. These protrusions could be made up of proteins with specific functions, e. g. facilitating membrane fusion (44) which could allow cargo delivery into the cytoplasm of a potential target cell. Similarly, only small double vesicles were found to contain a round incomplete structure which is thinner than a membrane bilayer. The nature of this structure is unknown but it could be a specific lumenal cargo. Both the surface coat and this cargo offered evidence of not only morphological but also biochemical differences in the isolated exosomes. A classification of vesicles purely based on their origin (apoptotic bodies, microvesicles and exosomes) is therefore insufficient to distinguish between different vesicle types. We classified exosomes derived from HMC-1 cell cultures into nine different categories based on their morphology.

Purification methods and sample preparation techniques may indirectly select for some subpopulations of vesicles with specific biochemical or physical characteristics over others, possibly affecting the ultimate outcome of experiments (45). Therefore, additional vesicle types that were not included in this study due to the chosen purification technique may exist. However, the observed categories are unlikely to be the result of an artifact caused by purification procedures, since very similar vesicle categories were also described for EVs in unprocessed ejaculate samples (15).

Vesicles within MVBs in HMC-1 cells showed a heterogeneous morphology as well, supporting our finding that exosomes are also structurally diverse. A large spectrum of morphologies was not limited to exosomes, as they were also observed in sectioned EVs found in cell culture (Figure 1), indicating that morphological variability is not uncommon among EVs.

Exosomes with non-spherical shapes and without any apparent scaffolding structure were observed, as in samples from other reports (11–13, 15, 16). It can be hypothesized that membrane-associated proteins, such as BAR-domain proteins or integral amphipathic helices, or the phospholipid composition itself may have a role in shaping the membrane of these vesicles (46–48). In other cases, filaments were observed in the lumen of vesicles and they were oriented in parallel to one another, possibly functioning as scaffolding for the shape of some vesicles (11, 12, 15). However, it is interesting to note that filaments were also found inside round vesicles, which is unprecedented. One hypothesis is that these vesicles were once tubular, but have lost their elongated shape due to a depolymerization of their internal filaments.

In conclusion, this study has showed a high degree of morphological variability in exosomes purified from the single cell type HMC-1. Different subpopulations of exosomes performing specific functions can therefore also be expected. We believe that this study will offer a better understanding of the nature of exosomes and EVs in general, which in turn could improve our general knowledge of cellular communication.

## Acknowledgements

We thank Bengt R Johansson and Richard Neutze for helpful discussions. We acknowledge the use of equipment at the Electron Tomography Facility at Karolinska Institutet and thank Sergej Masich for his help. JLH was supported by a VR young investigator grant and the Göran Gustafsson Foundation for Research in Natural Sciences and Medicine.

## Author contributions

J.L. and J.L.H designed research. A.C. isolated vesicles. J.L.H. did thin section EM and negative stain. C.L., M.B. and J.L.H. did cryo-electron microscopy. D.Z. analyzed the data. D.Z. and J.L.H. wrote the manuscript. All authors have given approval to the final version of the manuscript.

## Supplementary material

Figure S1: Complex structure of an agglomerate of vesicles. A-B) Cryo-electron micrograph of an agglomerate of vesicles and other electron dense material with drawing to illustrate its structure.

Table S1: Number of vesicles for each category and number of feature-presenting vesicles.

